# CRAIC: interactive interrogation of multiple sequence alignment uncertainty

**DOI:** 10.64898/2026.10.01.756022

**Authors:** James O. McInerney

## Abstract

Multiple sequence alignments (MSAs) are often treated as a solved problem. However, ambiguously or incorrectly aligned regions are the source of much error in phylogenetics and comparative genomics. CRAIC is a cross-platform desktop workbench that treats an alignment as a set of hypotheses to be stress-tested. CRAIC uses the posterior match probabilities of a three-state pair hidden Markov model (pair-HMM) to derive complementary views of alignment confidence. The five views are (i) a reliability overlay with live column and residue masking; (ii) a multi-aligner disagreement map, (iii) a local realignment sandbox, (iv) an interactive posterior explorer, and, where the true alignment is known, (v) a per-residue view of correctly predicted homologies. When present, MAFFT, MUSCLE, ProbCons, Clustal Omega, ClustalW and PRANK are integrated in the tool. In benchmarks the reliability score can locate misaligned columns, but while CRAIC can implement automatic filtering, it is primarily built for inspection and curation.

## Introduction

Multiple sequence alignments are statements of homology between individual residues along DNA or protein sequences. Robust alignments underpin several research fields including phylogeny reconstruction, tests of selection and comparative genomics (Ogden and Rosenberg, 2006; Wong et al., 2008; Fletcher and Yang, 2010). However, most alignments contain regions where assertions of homology are difficult to make, and these regions can disproportionately mislead downstream inference. Column-reliability approaches such as those implemented in the software program GUIDANCE and Heads-or-Tails use alignment perturbation approaches (Penn et al., 2010; Sela et al., 2015; Landan and Graur, 2007), while T-Coffee’s TCS and ZORRO use scores of internal alignment consistency (Chang et al., 2014; Wu et al., 2012). Both kinds of approach attempt to quantify, on a per-site basis, the confidence that can be placed in the robustness of the alignment. However, these methods typically produce either a static score related to alignment quality, or a trimmed alignment, and while this provides a principled approach to discarding some or all the alignment, it is possible to dive deeper into the problem. Jalview (Waterhouse et al., 2009) and AliView (Larsson, 2014) are general-purpose editors that provide excellent interactive views of an alignment. Since 2012, imported T-Coffee reliability scores can be used by Jalview to colour an alignment. However, that imported score can be neither interrogated nor recomputed when the alignment is edited, because it is employed as a static overlay that relates to only one specific alignment. The user is left with the choice of either working with a static filter or eyeballing the alignment without any assistance. The motivation for developing CRAIC was to close this gap by interrogating several complementary uncertainty signals, all derived from one probabilistic model, and providing them in a single program that can dynamically evaluate alternative alignments. Where the true alignment is known, CRAIC also shows exactly where each aligner goes wrong, which makes it a teaching tool as well.

### Features

CRAIC assumes a three-state (“match”, “insert”, and “delete”) pair-HMM with affine gaps. The method uses the idea of the posterior probability, denoted P(i∼j), that residue i of one sequence is homologous to residue j of a second sequence. The procedure calculates all these probabilities before producing the alignment itself. We use the ProbCons forward-backward procedure in log-space (Do et al., 2005). Figure 1 shows how all the core features in CRAIC emanate from this matrix. For speed and efficiency, the inner loop was programmed in the Rust language (via PyO3), but there is also a NumPy alternative implemented, and the program has the facility to fall back on it if the Rust core is not available.

**Figure 1.**
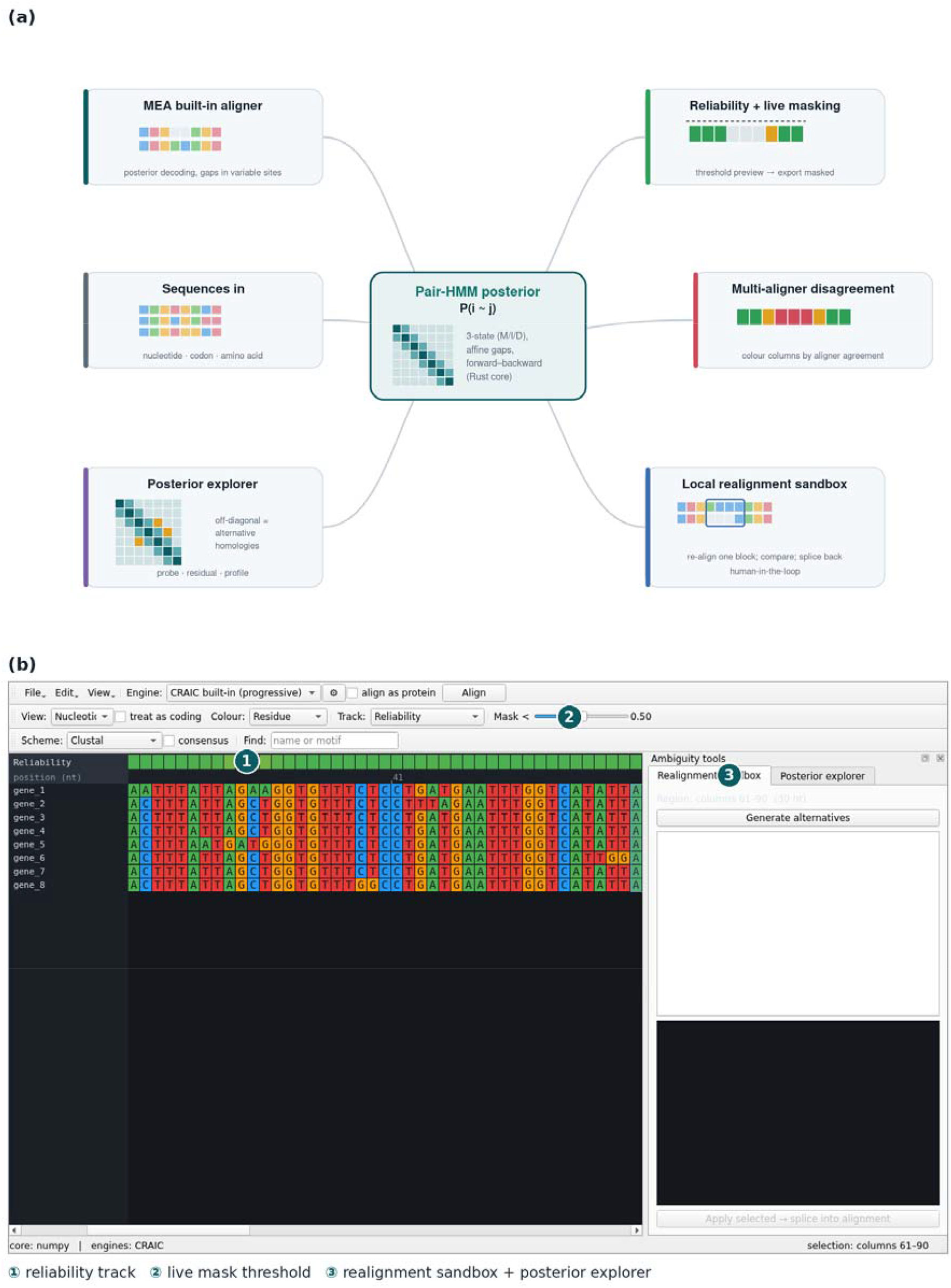
(a) A single three-state pair-HMM posterior matrix P(i∼j) (centre) drives P(i∼j) the built-in maximum-expected-accuracy aligner and all four uncertainty views: the reliability overlay with live masking, the multi-aligner disagreement map, the local realignment sandbox, and the posterior explorer. (b) The CRAIC interface on the bundled coding-gene example, showing □ the per-residue reliability track, □ the live mask-threshold slider (columns below the threshold are previewed before export), and □ the ambiguity-tools dock (realignment sandbox and posterior explorer), over an alignment shown at nucleotide level. Sequences are held in one canonical representation and can be shown at nucleotide, codon or amino-acid level.

Alignment confidence can be assessed in several ways, and we have implemented two main approaches in CRAIC to provide each column and residue with a confidence score. The first of these signals involves the computation of a consistency score like the implementation in TCS and ZORRO. The consistency score for a column is calculated as the mean posterior value over all the residue pairs in that column. The second signal is a stability probe that uses a perturbation approach like that used in GUIDANCE (Penn et al., 2010; Sela et al., 2015). CRAIC has a perturbation score implemented and it is calculated as the fraction of a residue’s homologies that survive being re-aligned using different approaches and different guide trees and gap regimes. The user can use a slider to preview which parts of the alignment would be included or excluded at any given threshold, before making the decision on a preferred alignment to export. Individual residues can also be masked, by hand or below a per-residue threshold, alone or together with the column mask. Every masked residue is then exported using ambiguity codes (N or X).

To facilitate filtering of the alignments, CRAIC implements several previously published approaches. Gap fraction trimming of alignments can be carried out using the trimAl approach with automatic gappyout trimming (Capella-Gutiérrez et al., 2009), as well as the Gblocks style (Castresana, 2000) of trimming conserved blocks of sequence. There is also the option of flagging an entire sequence that is in poor agreement with the rest of the alignment, using the OD-seq approach (Jehl et al., 2015). These options permit comparison of model-based and rule-based analyses.

Alignment ambiguity is most often investigated through the analysis of disagreement between independent alignment methods. CRAIC can be used to re-align sequences using all the available alignment engines (MAFFT, MUSCLE, Clustal Omega, ProbCons, PRANK and ClustalW, each with its own settings exposed), or if none are installed, the built-in aligner can be used with different gap regimes. These alignments can then be projected onto a common coordinate using shared residue identity. CRAIC then applies a colour to each column according to how often the different methods or regimes agree. We have also implemented a re-alignment sandbox. In this sandbox, the user can select a particular block of an existing alignment and ask for a re-alignment of this region using the various aligners that are available or using different gap costs and guide orders. The alternatives can then be compared side-by-side, and the best alignment can be manually selected and spliced back into the original, thereby replacing the region, but not disturbing the rest of the alignment. This step provides human-in-the-loop semi-manual curation of alignments.

Rather than displaying alignment columns as a single fixed alignment, the posterior explorer shows the user a local alignment as a heatmap, that represents the forward–backward posterior. In order to ask whether a residue could plausibly align in some other way, the user has the facility to select a variety of viewing options, including colour-by-confidence, per-residue homology-cloud probing, a residual mode that subtracts the committed alignment, and a one-versus-the-rest profile mode.

To demonstrate the difficulty of alignment, the program can show visually the relationship between alignment stability and correctness. This functionality is implemented in part to assist teaching in alignments, but also to understand the properties of the alignment approaches. Some of this idea was implemented in SuiteMSA Java program (Anderson et al., 2011). CRAIC can evolve sequences along a random tree, and the unaligned sequences can then be loaded into the graphical interface. The true alignment is known exactly but can be hidden from the user. The user can then align the sequences manually, or using any of the implemented automatic methods, and it dynamically reports which columns, and which individual residues are misplaced at any given time, next to the reliability score for those same columns. CRAIC will continue to report alignment correctness as the alignment is edited. A column inspector provides for every column information on which residues the reference groups there, those of them that ended up elsewhere, and any that do not belong. The user can also display the reference data itself, as a read-only alignment. A trusted structural reference such as BAliBASE can be used instead of the simulation, so the same tools can be used on real data. By providing the user with a comparison tool, as well as a statistical estimate of uncertainty and correctness, the user can check the alignment estimate against the errors themselves. ClustalW (Larkin et al., 2007) is included for the same purpose: its gap-opening and gap-extension costs can be changed and the consequences watched against the truth.

CRAIC has a built-in progressive aligner that estimates gap parameters from the data, and for nucleotide sequences, the emission divergence as well. For amino acid sequences an empirical substitution matrix is used. The idea is to use the ProbCons consistency transformation, which re-calculates every pairwise posterior using every third sequence as an intermediate. This means that evidence from all the sequences being aligned can be used to align every pair of residues. The process of decoding the model is where the alignment is produced, and the emitted alignment maximises the expected number of correctly aligned residue pairs, where the expectation is computed over the posterior itself. Gap opening and extension costs are already embedded in the pair-HMM’s transition parameters, and these are estimated from the sequences unless the user sets them.

This built-in progressive aligner is intended to be always available as a relatively fast alignment tool and is not expected to out-perform dedicated tools such as MAFFT (Katoh and Standley, 2013), MUSCLE (Edgar, 2004; 2022) or Clustal Omega (Sievers et al., 2011; Table 1). Protein-coding regions that are input in nucleotide format can be aligned in amino-acid space and back-translated, in order to ensure that indels are grouped into threes, and consequently respect the reading frame, in the spirit of MACSE (Ranwez et al., 2011) and phylogeny-aware placement (Löytynoja and Goldman, 2008). The canonical representation projects instantly between nucleotide, codon and amino-acid views.

**Table 1.** Alignment accuracy and reliability, by aligner. Sum-of-pairs (SP) is recall, the fraction of the reference’s homologous residue pairs recovered. Total-column (TC) is the fraction of reference columns reproduced exactly. AUC is the ROC area of a reliability score predicting which columns are correct, computed with midranks so that heavily tied scores do not make the answer depend on input order. BAliBASE SP and TC are core-block scores from the official bali_score. Simulation figures are means over 360 datasets. Every aligner was run with its own default settings, except that CRAIC’s built-in engine was run at its highest compute level on BAliBASE. Full tables, with standard errors and per-set and per-divergence breakdowns, are given in Supplementary Methods.

|  | BALiBASE 3 (163 families) |  |  | Simulation (360 datasets) |  |
| --- | --- | --- | --- | --- | --- |
| Aligner | SP | TC | AUC | SP | AUC |
| **CRAIC built-in** | 0.830 | 0.676 | 0.848 | 0.604 | 0.855 |
| **MAFFT 7.505** | 0.818 | 0.667 | 0.874 | 0.608 | 0.893 |
| **MUSCLE 5.1** | 0.856 | 0.712 | 0.833 | 0.663 | 0.858 |
| **Clustal Omega 1.2.4** | 0.766 | 0.593 | 0.876 | 0.549 | 0.943 |
| **ProbCons 1.12** | 0.850 | 0.705 | 0.845 | - | - |
| **PRANK v.170427** | 0.714 | 0.506 | 0.905 | - | - |
| **ClustalW 2.1** | 0.743 | 0.551 | 0.903 | 0.555 | 0.927 |

### Example and performance

The Supplementary Methods provide the full protocol and per-set results. On real data we used BAliBASE 3 (Thompson et al., 2005), re-aligning each family with the built-in engine and with MAFFT 7.505, MUSCLE 5.1, Clustal Omega 1.2.4, ProbCons 1.12, PRANK v.170427 and ClustalW 2.1. The analysis was restricted to the 163 families that every aligner completed (reference sets RV11 in full, and all but one family of RV12; the consistency transformation is cubic in the number of sequences and does not scale to the larger sets, which were not attempted), and scoring was carried out using the official bali_score over the reference core blocks. On simulated data we used 360 datasets of either 8 or 16 sequences, with a known-true alignment and tree, spanning six divergence levels from 0.1 substitutions per site to 1.5 subs/site, and we used a sweep of three indel rates.

Table 1 summarises both the real data and the simulations. The reliability score identifies mis-aligned columns, with a ROC AUC of 0.83–0.90 on real alignments and 0.86–0.94 on our simulated data. This shows that the reliability score performs at least as well on alignments CRAIC did not build as on those it did. Furthermore, unlike alignment accuracy itself, reliability does not degrade with divergence, and the reliability scores remained between 0.83 and 0.89 across the simulated range while, in contrast, sum-of-pairs accuracy falls from 0.98 to 0.19 (see Supplementary Information). Finally, when we masked the flagged columns the mean per-column accuracy for the retained fraction of the alignments was between 0.819–0.858 on BAliBASE, compared with accuracy scores of 0.592–0.671 for masks that discarded the same number of columns at random and 0.669–0.733 for gap-fraction masks of the same size (z = +8.0 to +11.3).

Masking improves per-column accuracy, but its effect on trees depends on how much it removes (figure 2). At 20–40% identity (BAliBASE RV12) the mask removed a median of 28% of columns and left trees from MAFFT alignments as close to the tree of the reference alignment as before (Robinson–Foulds distance 0.26 against 0.27, p = 0.23), whereas removing as many columns at random did not (0.32). Below 20% identity (RV11) it removed a median of 83%, and the resulting trees were less accurate (0.55 against 0.39, p = 0.004). Heavy masking discards phylogenetic signal along with ambiguity, consistent with reports that automated alignment filtering can worsen phylogenetic inference (Tan et al., 2015). Masking residues instead, which is reported to lose less signal (Ali et al., 2019; Wheeler et al., 2026), did not recover the loss (figure 2). The reliability signal is thus best used to locate and interrogate ambiguous regions, not as an automatic filter. All analyses are interactive on a laptop with the compiled core.

**Figure 2.**
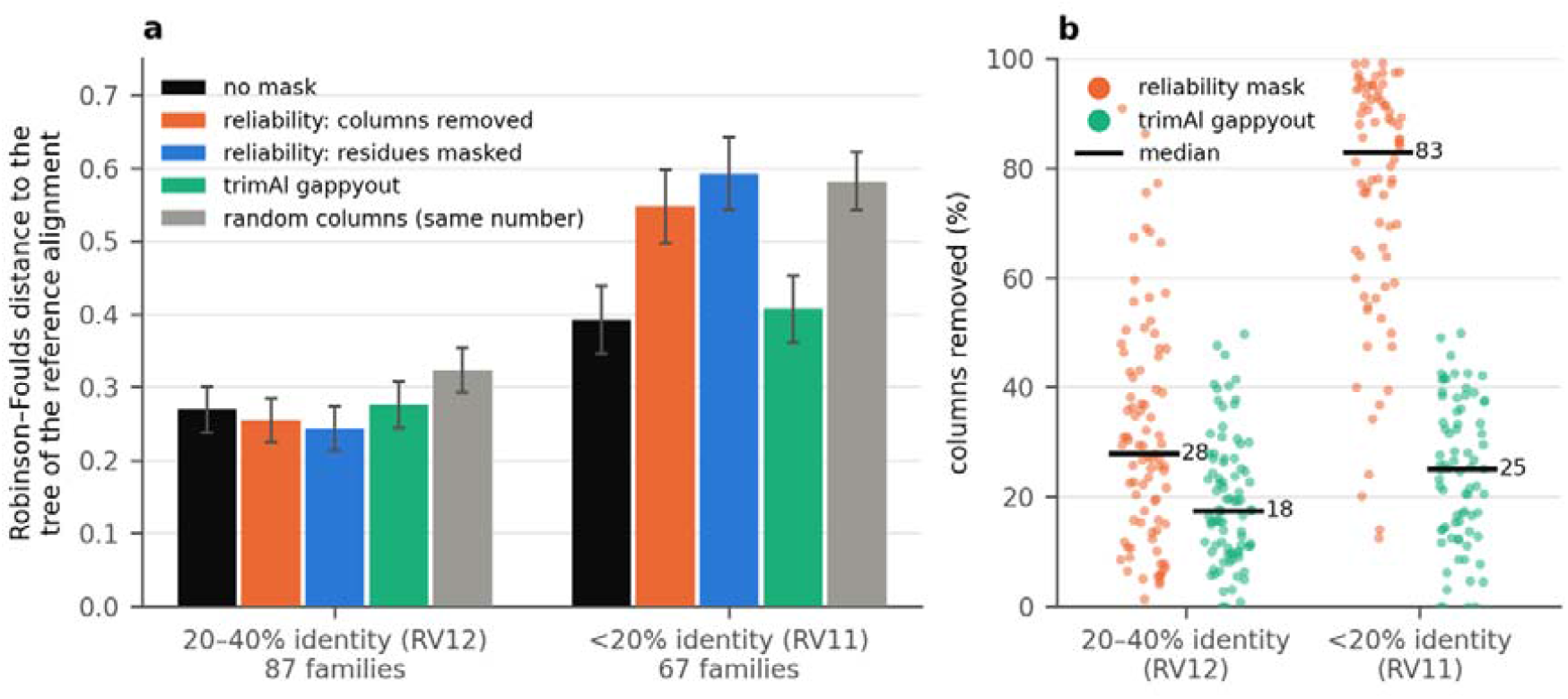
On real alignments, masking by reliability harms trees only when it removes most of the alignment. (a) Normalised Robinson–Foulds distance (Robinson and Foulds, 1981) between the maximum-likelihood tree (IQ-TREE 2, LG+G4; Minh et al., 2020) of each MAFFT alignment of a BAliBASE family and the tree of its reference alignment: unmasked; with columns scoring below 0.5 removed; with residues scoring below 0.5 masked as missing data; after trimAl gappyout; and with as many columns as the reliability mask removed, chosen at random (mean of three draws). Means ± s.e. over the families in which every mask left enough data for a tree. (b) Share of each family’s columns removed by the reliability mask and by gappyout.

## Conclusion

CRAIC has been designed to reframe the multiple sequence alignment from being a fixed input for downstream analyses, into an object of active real-time interrogation. CRAIC gives the curator several independent, model-based views of where an alignment can and cannot be trusted. It also provides the tools to act on that judgement, before the alignment is passed to phylogenetic or comparative analysis. The exploration of alignment uncertainty becomes a routine part of curating an alignment, unifying reliability scoring, multi-method disagreement, local realignment, residue masking and posterior exploration on one pair-HMM primitive in a single interactive application.

## Supplementary Material

Supplementary Methods give the full specification of the model, the estimators and the reliability ensembles, with per-set and per-divergence results. Documentation and a guided tutorial are at https://mol-evol.github.io/craic/.

## Data Availability

CRAIC is available as free, open-source software under the MIT licence at https://github.com/mol-evol/craic and on PyPI (*pip install craic-msa*); standalone desktop bundles for macOS, Windows and Linux are provided. It is implemented in Python (PySide6) with a compiled Rust core and a cross-validated NumPy fallback and requires Python 3.9+. The benchmark scripts, simulation settings and raw results behind Table 1 and figure 2 are in the same repository (benchmarks/).

## Supplementary Methods

### CRAIC: interactive interrogation of multiple sequence alignment uncertainty

James O. McInerney

The matrix of posterior match probabilities for a pair of sequences forms the basis for most calculations in CRAIC. This document provides an outline of (i) how this matrix is constructed, (ii) the way in which its parameters are estimated, (iii) how it is then decoded into an alignment, and (iv) the way it is used as a measure of reliability. A version of this material that will be continuously updated can be found at https://mol-evol.github.io/craic/methods/.

### S1 The pair-HMM

Two ungapped sequences a (length m) and b (length n) are said to be related by a three-state hidden Markov model, with states M, X and Y. M is the state in which the two residues are homologous, X is the state in which a residue of a is aligned to a gap, and Y is the state in which a residue of b is aligned to a gap. Gaps are affine, and we can parameterize them by two numbers. The first is δ, which is the probability of changing from state M for X or Y (gap opening), and the second is ε, which is the probability of remaining in state X or state Y (gap extension). The transitions are M→M = 1 − 2δ, M→X = M→Y = δ, X→X = Y→Y = ε and X→M = Y→M = 1 − ε. This is the formulation that used in the ProbCons software (Do et al., 2005).

State M emits a residue pair from a joint distribution J; X and Y emit a single residue from the background distribution p. For nucleotide sequences, we build J from an θ which is an estimated identity, with diagonal mass θ/4, off-diagonal mass (1 − θ)/12 and we use a uniform background of 0.25. For proteins, J is already built and is simply the BLOSUM45 matrix (which, in our hands, scored better than the more usual BLOSUM62 on BAliBASE) converted to a proper joint. BLOSUM45 entries are third-bit log-odds, 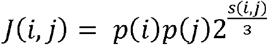, normalised to sum to one, with p being the Robinson and Robinson background frequencies. The user is presented with a dropdown menu so they can select between alternative substitution matrices at runtime. The BLOSUM45 matrix is the default matrix, but other options include BLOSUM50, 62, 80 or 90. This does carry the consequence that the same model is used irrespective of whether the sequences are 80% identical or whether they the sequences are 25% identical.

#### S1.1 Parameter estimation

A committed alignment is turned into pair-HMM parameters by a single function, and is it then used by both the aligner’s own pass and by every reliability score. From the alignment the mean pairwise identity over aligned residue pairs, θ, is calculated and this is clamped to [0.55, 0.95], which we feel are sensible safe limits. Also δ, gap openings divided by the sum of residue positions and gap openings, is calculated and it is clamped to [0.005, 0.2]. Finally, ε is calculated, which is one minus the ratio of openings to gapped positions, and this is clamped to [0.1, 0.9]. In the case where there is no alignment to estimate from, the defaults for the library (δ = 0.02, ε = 0.5) will apply.

#### S1.2 Forward–backward

In order to calculate the posterior, we use forward–backward in log space. The resulting matrix, P(i∼j), is the probability that residue i of a is homologous to residue j of b, and this is marginalised over every alignment of the pair, and each one of these is weighted by its probability under the model. CRAIC treats this matrix, rather than any single alignment, as the state of knowledge about the pair.

### S2 The consistency transformation

The ProbCons consistency transformation re-estimates each posterior using every third sequence: P′(x,y) = (1/n) Σ_z P(x,z)P(z,y) and in that way propagates evidence through the family. We apply the transformation for two iterations by default. Where x and y both align a residue of z to each other’s residue, that agreement is what raises the posterior even when the direct evidence is weak.

The consistency transformation is the most expensive operation that is performed by the CRAIC tool. Performance grows proportionately to the cube of the input size, known as “Big O cubed” (O(n^3^) products of L×L matrices per iteration). This places a limit on the largest family the workbench can handle. Most of these probabilities end up being close to zero, so just like ProbCons, CRAIC can discard those below 0.01 and store only the rest, and this typically reduces the requirement to 1.8–6% of the memory. This is, of course, an approximation, and it has the implication that on divergent sequences, many of the probabilities are very low. In fact, on the simulated DNA alignments, with divergence set to 0.5 substitutions per site, the built-in engine recorded a reduction of approximately 0.1 of sum-of-pairs accuracy. CRAIC therefore uses the exact transformation whenever it fits in the memory budget, and it uses the sparse option only when memory requirements become excessive for the computer on which it is running, and if SciPy is installed.

### S3 Building an alignment S3.1 Guide tree

Guide-tree distances are obtained using the posteriors that were computed earlier. The distance d(x,y) is calculated as d(x,y) = 1 − min(1, ΣP(x,y) / min(|x|,|y|)), that is, one minus the expected number of aligned residues over the length of the shorter sequence. If the lowest compute level is being used, where no all-pairs posteriors are retained in memory, the tree falls back to using the cosine distance on 3-mer counts. Sequences are merged by nearest-neighbour joining with proportional averaging of distances to the merged profile.

#### S3.2 Maximum-expected-accuracy decoding

As in ProbCons (Do et al. 2005), CRAIC uses a method of decoding by maximum expected accuracy. This means that over all alignments A, it maximises Σ P(i∼j) for (i,j) in A, using dynamic programming over the posterior matrix. There is no gap penalty; the cost of a gap is already accounted for in the posterior as δ and ε. At each node of the guide tree, the two groups of already-aligned sequences are joined together, and this relative alignment is fixed going in to the next round. This joining uses the same criterion, using the posteriors summed over the residue pairs that would be aligned by the joining.

#### S3.3 Scale, and codon-aware alignment

Holding every pairwise posterior is O(n^2^L^2^) in memory, so a doubling of the sequence length or a doubling of the number of sequences will quadruple memory usage.

Above a stated budget the aligner computes each posterior on demand and discards it. This means that peak memory is a single matrix rather than n^2^. This allows the program to scale up reasonably well, though it relinquishes some accuracy in the consistency transformation. In the event that memory is a limitation, this trade-off is reported to the user. CRAIC also implements a codon-aware alignment function, where the sequences are dynamically translated (using the selected genetic code) into amino acid sequences, aligned at the protein level and then back-translated using their original DNA sequences. This means the alignment can be represented as an alignment of codons, which is useful for analyses of selection in particular.

### S4 Reliability

We calculate the consistency score by taking every column and computing the mean posterior over each of the residue pairs contained in that column. For very large families we restrict the calculation to a maximum of 300 sequence pairs. A column will be given a low score if it has grouped together a set of residues that the pair-HMM considers unlikely to be homologous. This procedure was developed in the spirit of TCS from the T-Coffee program and in the spirit of ZORRO but the difference being that it is computed over CRAIC’s own posteriors. This means the numbers are not interchangeable. Because of how it works, consistency can be reported on a per column, and on a per residue basis. A residue that has no partners in its column is left undefined.

We calculate the perturbation score by re-aligning the same sequences sixteen times with perturbed guide trees, while also cycling through three gap regimes (δ = 0.01, 0.03, 0.08). Then, for each residue, we then score the fraction of the originally asserted homologies that survive this ensemble of perturbations. Guide trees are perturbed by bootstrap-resampling the k-mer feature columns, and the program uses a light jitter in order to resolve near-ties. In contrast, GUIDANCE uses a procedure where it bootstraps alignment columns, typically over 100 replicates, and from there it rebuilds guide trees and re-runs the aligner that produced the reference.

Reliability can be computed as the average of the consistency and perturbation scores. Columns can be masked and exported based on these scores.

### S5 Benchmark protocol

We benchmarked the alignment methods using two independent sources of ground truth and the results were scored in two different ways. The sum-of-pairs (SP) metric is calculated as the fraction of the reference’s homologous residue pairs that were recovered. This is often called recall in statistical analysis. The total-column (TC) metric is the fraction of reference columns reproduced exactly. The per-column correctness is the fraction of a column’s asserted pairs that are true, and we set this as undefined in the event that a column asserts none. Finally, reliability AUC is the ROC area of a reliability score predicting which columns are correct. Reliability AUC is computed with midranks so that heavily tied scores are not overly-influences by input order.

The real data that we used came from the BAliBASE 3 database. The sequences were re-aligned with the built-in engine and with MAFFT 7.505, MUSCLE 5.1, Clustal Omega 1.2.4, ProbCons 1.12, PRANK v.170427 and ClustalW 2.1. These alignments were then scored using the official bali_score over the reference core blocks and, separately, with CRAIC’s own all-column metrics. The analysis covers the 163 families of reference sets RV11 (76/76) and RV12 (87/88). One family was excluded from analysis because its dense pairwise posteriors exceeded the 3 GB memory budget. RV20–RV50 were not attempted for this analysis due to the excessive memory requirements of such large datasets. Every alignment program was run with its own default settings, except that the built-in engine was run at its highest compute level (max; Table S1) here and at its default (med) in the simulations.

The simulated data used 360 datasets that we evolved down random binary trees with substitutions, insertions and deletions and gamma-distributed among-site rate variation (α = 0.8), over 8 and 16 taxa, root length 400 nt, six divergence levels and three indel rates, with 10 replicates per condition.

### S6 Results

**Table S1.** Compute levels. The consistency transformation is what distinguishes them.

| Compute level | Consistency iterations | Guide tree | Refinement |
| --- | --- | --- | --- |
| min | 0 | k-mer | 0 |
| med (default) | 2 | posterior | 0 |
| max | 2 | posterior | 25 |

**Table S2.** BAliBASE 3, official core-block bali_score, 163 families. Mean ± standard error.

| Aligner | SP | TC |
| --- | --- | --- |
| MUSCLE 5.1 | 0.856 ± 0.012 | 0.712 ± 0.020 |
| ProbCons 1.12 | 0.850 ± 0.013 | 0.705 ± 0.021 |
| CRAIC built-in | 0.830 ± 0.014 | 0.676 ± 0.022 |
| MAFFT 7.505 | 0.818 ± 0.016 | 0.667 ± 0.022 |
| Clustal Omega 1.2.4 | 0.766 ± 0.018 | 0.593 ± 0.024 |
| ClustalW 2.1 | 0.743 ± 0.017 | 0.551 ± 0.025 |
| PRANK v.170427 | 0.714 ± 0.018 | 0.506 ± 0.023 |

**Table S3.** BAliBASE 3, CRAIC’s all-column scoring, which the per-column analyses require. The reliability AUC is at least as high on alignments CRAIC did not build as on its own.

| Aligner | SP | TC | Reliability AUC |
| --- | --- | --- | --- |
| MUSCLE | 0.740 | 0.466 | 0.833 ± 0.010 |
| ProbCons | 0.732 | 0.460 | 0.845 ± 0.009 |
| CRAIC built-in | 0.714 | 0.445 | 0.848 ± 0.007 |
| MAFFT | 0.700 | 0.431 | 0.874 ± 0.007 |
| Clustal Omega | 0.665 | 0.399 | 0.876 ± 0.009 |
| ClustalW | 0.643 | 0.379 | 0.903 ± 0.009 |
| PRANK | 0.596 | 0.332 | 0.905 ± 0.007 |

**Table S4.**
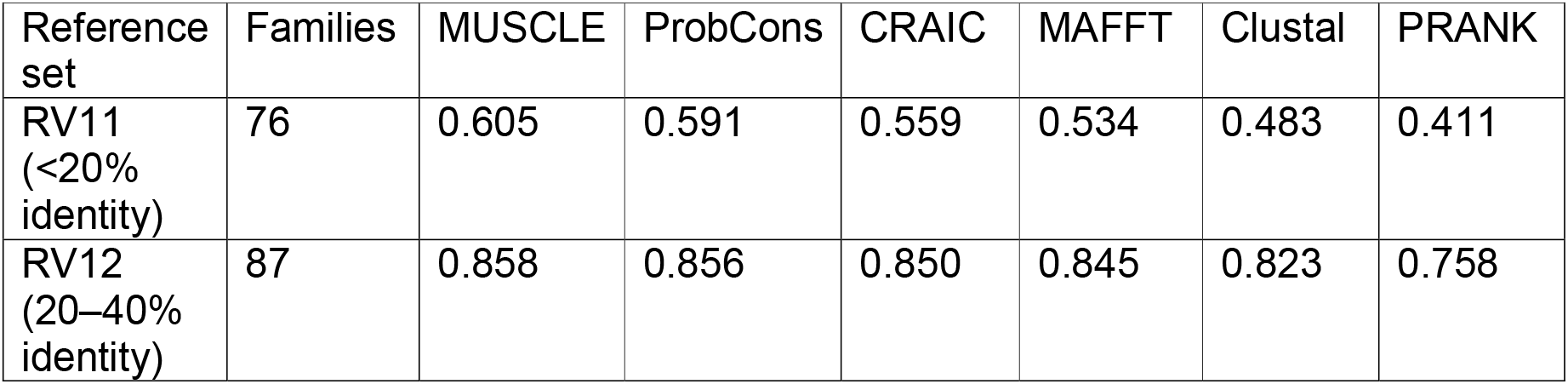
BAliBASE SP by reference set, all-column scoring. Below 20% identity every method loses roughly half the true homologies, which is the regime in which knowing which columns to distrust matters most. ClustalW: 0.468 (RV11) and 0.797 (RV12).

**Table S5.** Masking against controls, BAliBASE. Mean per-column accuracy of the retained alignment, against a mask discarding the same number of columns at random and against a gap-fraction mask of the same size. The margin over the gap rule is the informative one.

| Aligner | Kept columns | Matched random | Gap-fraction rule | z |
| --- | --- | --- | --- | --- |
| ClustalW | 0.858 | 0.606 | 0.669 | +11.3 |
| PRANK | 0.846 | 0.592 | 0.671 | +10.8 |
| Clustal Omega | 0.842 | 0.615 | 0.710 | +9.8 |
| MUSCLE | 0.837 | 0.671 | 0.733 | +8.1 |
| ProbCons | 0.832 | 0.666 | 0.733 | +8.1 |
| CRAIC built-in | 0.819 | 0.648 | 0.714 | +8.0 |
| MAFFT | 0.828 | 0.634 | 0.728 | +9.5 |

**Table S6.** Simulation, 360 datasets, means over all conditions. The reliability AUC is higher here than on BAliBASE because the simulated sequences are generated under a model closer to the pair-HMM that computes the score; the BAliBASE figures are the conservative ones.

| Aligner | SP | TC | AUC | Kept | Random | Gap rule | z |
| --- | --- | --- | --- | --- | --- | --- | --- |
| CRAIC built-in | 0.604 | 0.369 | 0.855 | 0.926 | 0.529 | 0.690 | +6.9 |
| MAFFT | 0.608 | 0.379 | 0.893 | 0.843 | 0.533 | 0.548 | +11.3 |
| MUSCLE | 0.663 | 0.387 | 0.858 | 0.884 | 0.581 | 0.669 | +8.5 |
| Clustal Omega | 0.549 | 0.261 | 0.943 | 0.836 | 0.529 | 0.532 | +12.2 |
| ClustalW | 0.555 | 0.332 | 0.927 | 0.873 | 0.534 | 0.534 | +12.6 |

**Table S7.** Simulation by divergence. Columns 2–5 are SP. RF is the Robinson– Foulds distance of the neighbour-joining tree to the true topology, for CRAIC alignments, unmasked and masked at the default 0.5 threshold. Accuracy falls off a cliff between 0.25 and 0.5 for every method, while the reliability AUC on CRAIC alignments holds between 0.83 and 0.89 across the whole range. ClustalW SP by divergence: 0.955, 0.825, 0.601, 0.435, 0.333, 0.180.

| Divergence | CRAIC | MAFFT | MUSCLE | Clustal Omega | RF full | RF masked |
| --- | --- | --- | --- | --- | --- | --- |
| 0.1 | 0.982 | 0.965 | 0.964 | 0.860 | 0.00 | 0.00 |
| 0.25 | 0.914 | 0.887 | 0.892 | 0.746 | 0.04 | 0.14 |
| 0.5 | 0.697 | 0.680 | 0.730 | 0.594 | 0.16 | 0.42 |
| 0.75 | 0.484 | 0.509 | 0.590 | 0.473 | 0.29 | 0.60 |
| 1.0 | 0.354 | 0.396 | 0.490 | 0.382 | 0.42 | 0.68 |
| 1.5 | 0.191 | 0.214 | 0.314 | 0.241 | 0.60 | 0.81 |

Masking improves per-column accuracy substantially and consistently, and makes neighbor-joining tree recovery worse at every divergence above the trivial. At high divergence the threshold discards most of the columns, and this eliminates most of the phylogenetic signal. To put it another way: a more accurate alignment that only includes a quarter of the data is a worse basis for phylogenetic tree reconstruction than a less accurate alignment of all of the data. This is consistent with Tan et al. (2015). The reliability signal should really be used to locate and interrogate ambiguous regions, and to curate them by hand. It should not be used as an automatic filter as a prelude to a search for a phylogenetic tree.

S6.1 No method can reconstructs all of the alignments (or even very many of them) Across the same 163 families, seven aligners and the official core-block scorer, MUSCLE, ProbCons, ClustalW and the built-in engine each reconstruct 6 families exactly (total-column score 1.000), MAFFT 5, and Clustal Omega and PRANK none. Eleven families are reconstructed exactly by at least one method; not one family is reconstructed exactly by all seven. Nor is the best method knowable in advance: MUSCLE produces the best alignment for 81 of the 163 families, leaving 82 won by one of the other six, and the built-in engine is best for 32 of them. An oracle allowed to choose the best method per family would reach a total-column score of 0.742 against MUSCLE’s 0.712 — an improvement unavailable in practice that still leaves a quarter of columns misplaced. These are capable tools and each recovers the large majority of homologous pairs; the point is that no automatic procedure returns an alignment that can be treated as settled, and that the errors differ from method to method.

### S6.2 Reliability AUC is not a ranking of aligners

Table S3 must not be read as ordering the aligners by quality: across the seven engines, reliability AUC and alignment accuracy are perfectly anticorrelated (Spearman ρ = −1.00, p = 0.0004). The AUC is highest on PRANK’s alignments, which are the least accurate, and lowest on MUSCLE’s, which are the most. This is expected rather than perverse. A less accurate alignment contains more genuinely wrong columns, so there is more signal for any classifier to separate, while a nearly-correct alignment offers few errors to find. The column reports how well the score identifies wrong columns given an alignment; it carries no information about which alignment to prefer.

### S7 For reproduction

The software and the raw output of every run reported here is included under benchmarks/results/. Commands for reproducing the paper’s results are in docs/validation.md and at https://mol-evol.github.io/craic/validation/.

